# T cell stimulation remodels the latently HIV-1 infected cell population by differential activation of proviral chromatin

**DOI:** 10.1101/2021.10.04.461021

**Authors:** Birgitta Lindqvist, Wlaa Assi, Julie Roux, Luca Love, Bianca B. Jütte, Anders Sönnerborg, Tugsan Tezil, Eric Verdin, J. Peter Svensson

## Abstract

The reservoir of latently HIV-1 infected cells is heterogeneous. To achieve an HIV-1 cure, the reservoir of activatable proviruses should be eliminated while permanently silenced proviruses may be tolerated. We have developed a method to assess the proviral nuclear microenvironment in single cells. In latently HIV-1 infected cells, a zinc finger protein tethered to the HIV-1 promoter produced a fluorescent signal as a protein of interest came in its proximity, such as the viral transactivator Tat when recruited to the nascent RNA. Tat is essential for viral replication. In these cells we assessed the proviral activation and chromatin composition. By linking Tat recruitment to proviral activity, we dissected the mechanisms of HIV-1 latency reversal and the consequences of HIV-1 production. A pulse of promoter-associated Tat was identified that contrasted to the continuous production of viral proteins. As expected, promoter H3K4me3 led to substantial expression of the provirus following T cell stimulation. However, the activation-induced cell cycle arrest and death led to a surviving cell fraction with proviruses encapsulated in repressive chromatin. Further, this cellular model was used to reveal mechanisms of action of small molecules. In a proof-of-concept study we determined the effect of an enhancer specific CBP/P300-inhibitor on HIV-1 latency reversal. Only proviruses resembling active enhancers, associated with H3K4me1 and H3K27ac, efficiently recruited Tat. Tat-independent HIV-1 latency reversal of unknown significance still occurred. We present a method for single cell assessment of the microenvironment of the latent HIV-1 proviruses, used here to reveal how T cell stimulation modulates the proviral activity and how the subsequent fate of the infected cell depends on the chromatin context.

## Introduction

Despite antiretroviral therapy (ART) potently inhibiting HIV-1 replication, the intact HIV-1 genome persists in infected cells. Most of these cells do not produce viral particles and hence make up the latent reservoir. Some proviral transcription is observed in these cells but rarely are the transcripts translated (Yukl et al., 2018). Thereby the cells evade immune recognition. Efforts to stimulate the cells and expose them as infected to the immune system has failed clinically as even the strongest activators lead to limited HIV-1 latency reversal (Battivelli et al., 2018; Sarabia et al., 2021). Among people living with HIV-1, rare individuals naturally control viral replication, a control driven by a potent immune response that kill cells exposing viral proteins (Migueles et al., 2008) in combination with a majority of proviruses integrated in repressive chromatin (Jiang et al., 2020). This combined approach of eliminating the reservoir of activatable proviruses while tolerating permanently silenced proviruses, produces a sustained, drug-free HIV-1 remission–a *de facto* HIV-1 cure.

To reproduce this HIV-1 remission clinically in a more general population, we must specifically target each sub-compartment of the latent reservoir subsequently. As discrete sub-compartments may be under specific control, first we need to delineate the distinct mechanisms governing HIV-1 latency reversal (Sherrill-Mix et al., 2013). A well-studied reservoir fraction contains proviruses integrated in active regions. Where the HIV-1 promoter is associated with the H3K4me3 chromatin mark. These open chromatin structures contain most proviruses upon initial integration (Chen et al., 2017; Battivelli et al., 2018). As the HIV-1 capsid enters the nucleus through the nuclear pore (Bejarano et al., 2019; Blanco-Rodriguez et al., 2020), the provirus almost immediately integrates, usually within a short distance (Marini et al., 2015; Burdick et al., 2020). The microenvironment of the nuclear pore is a highly transcriptionally permissive location (Capelson et al., 2010). Even intron-containing HIV-1 transcripts may be exported through the nuclear pore once chaperoned by the viral Rev protein (Coyle et al., 2011). Stochastic induction of latency is reversed upon T cell activation when growth factors and nutrient supply become abundant and the NFκB transcription factor binds to the promoter region. Metabolic activity enables HIV-1 latency reversal (Besnard et al., 2016). The gene-like structure allows mRNA processing and translation of viral proteins, including the essential transactivator of transcription Tat. Another reservoir sub-compartment is contained in enhancer-like structures. Also these proviruses remain accessible and with high reactivation potential (Chen et al., 2017; Battivelli et al., 2018; Lucic et al., 2019). Enhancer chromatin structures enable long-term reactivatable latency as it provides an open chromatin without mRNA production (Lindqvist et al., 2020). In contrast, the initially rare cellular fractions where proviral genomes are encapsulated in heterochromatin marks H3K9me3 or H3K27me3 have low reactivation potential (du Chene et al., 2007; Kauder et al., 2009; Battivelli et al., 2018). H3K9me3 contained proviruses may still be activated by knock-down of heterochromatin protein 1 (HP1) (du Chene et al., 2007) but their clinical contribution is questionable. Further, the HIV-1 reservoir is dynamic and evolves over time (Bruner et al., 2019; Pinzone et al., 2019; Simonetti et al., 2020). Proviruses in heterochromatin gradually take on a larger fraction of the HIV-1 infected cells, on expense of proviruses with components of actively transcribed regions (Lindqvist et al., 2020). Heterochromatin forms over the HIV-1 promoter in the absence of functional Tat (Li et al., 2019).

For *in vivo* replication of the HIV-1, the Tat protein is essential. In an early step of HIV-1 latency reversal, Tat is recruited to the promoter to override the host-controlled transcription machinery and augment proviral expression (Morton et al., 2019). Tat binds to an initial nucleotide sequence of the nascent HIV-1 RNA. It then recruits and modifies the P-TEFb complex to enable transcription elongation. Once host mechanisms are assembled for transcription, Tat is acetylated and loses its affinity for RNA (Kiernan et al., 1999; Kaehlcke et al., 2003). In the viral circuitry, Tat has been proposed to generate promoter toggling, *i*.*e*. being responsible for both the ON and OFF switch of the HIV-1 provirus (Razooky et al., 2017). Except for regulating the HIV-1 promoter, ectopic expression of Tat has been observed to have affinity for other genomic regions (Marban et al., 2011; Reeder et al., 2015). Apart from promoting transcription, Tat stimulates activation induced cell death (AICD) (Gülow et al., 2005). AICD is a naturally occurring process where activated T cells induce apoptosis after an immune response as to maintain the balance of T cells once infection is cleared. In this process, the cells progress through the cell cycle and arrest in G1 (Lissy et al., 1998). As cell stress and death in itself induce HIV-1 reactivation (Khan et al., 2015) and cells with activated HIV-1 express toxic viral proteins that further reduce the viability of virus, this generates a positive feed-back loop of HIV-1 latency reversal and cell death. This process becomes particularly strong in cellular models. In models with a late readout such as the accumulated production of p24 (or a reporter protein such as GFP), HIV-1 latency reversal induced by T cell activation or as a consequence of AICD, become inseparable.

Here we present a single cell method to assess chromatin at the HIV-1 locus and link it to early stages of HIV-1 activation. Using this method, we show that upon T cell stimulation, Tat was transiently recruited to the HIV-1 promoter in proliferating cells. Following T cell stimulation, the proviral chromatin landscape within the HIV-1 infected cells was altered due to selective cell death. We also found that Tat was uniquely present at the HIV-1 promoter when it had enhancer features delivered by CBP/P300.

## Results

### Labeling of the HIV-1 provirus to enable proximity ligation assay at the promoter

To study the microenvironment of the HIV-1 promoter in single cells, we adapted the well-established proximity ligation assay (PLA) (Soderberg et al., 2006). Performing PLA in fixed cells attached to microscope cover slips results in a bright fluorescent signal when two query proteins are within 40 nm of each other (Fig 1A). PLA relies on the proximity of two antibodies that have been conjugated with an oligonucleotide probe. To enable detection of the integrated HIV-1, we tethered a small protein to the proviral promoter region. Zinc finger proteins (ZFP) bind to DNA sequence in a highly specific manner. A specific protein, ZFP3, has been artificially constructed to recognize the HIV-1 promoter, adjacent to the NFκB binding sites (Wang et al, 2014). We cloned a FLAG-labelled ZFP3 protein into a lentiviral plasmid under the SFFV promoter. The ZFP3 sequence was linked to the BFP sequence through the sequence for a self-cleaving P2A peptide. Lentiviral particles were transduced into J-lat 5A8 cells, a Jurkat cell line harboring a single copy of latent HIV-1 (Jordan et al., 2003; Chan et al., 2013). The latent reporter HIV-1 in 5A8 cells is a full-length mutated provirus where *env* has a frame-shift mutation and *nef* is replaced by a GFP coding sequence. After clonal expansion of ZFP3-BFP transduced cells, for further studies we selected a clone, 1C10, with 96.4 % ZFP3 positive cells, as identified by BFP expression (Fig S1A). The FLAG-ZFP3 protein was present in 1C10 cells in both unstimulated cells and cells stimulated with phorbol 12-myristate 13-acetate and ionomycin (PMA/i) as confirmed by immunoblotting (Fig 1B).

**Fig 1.**
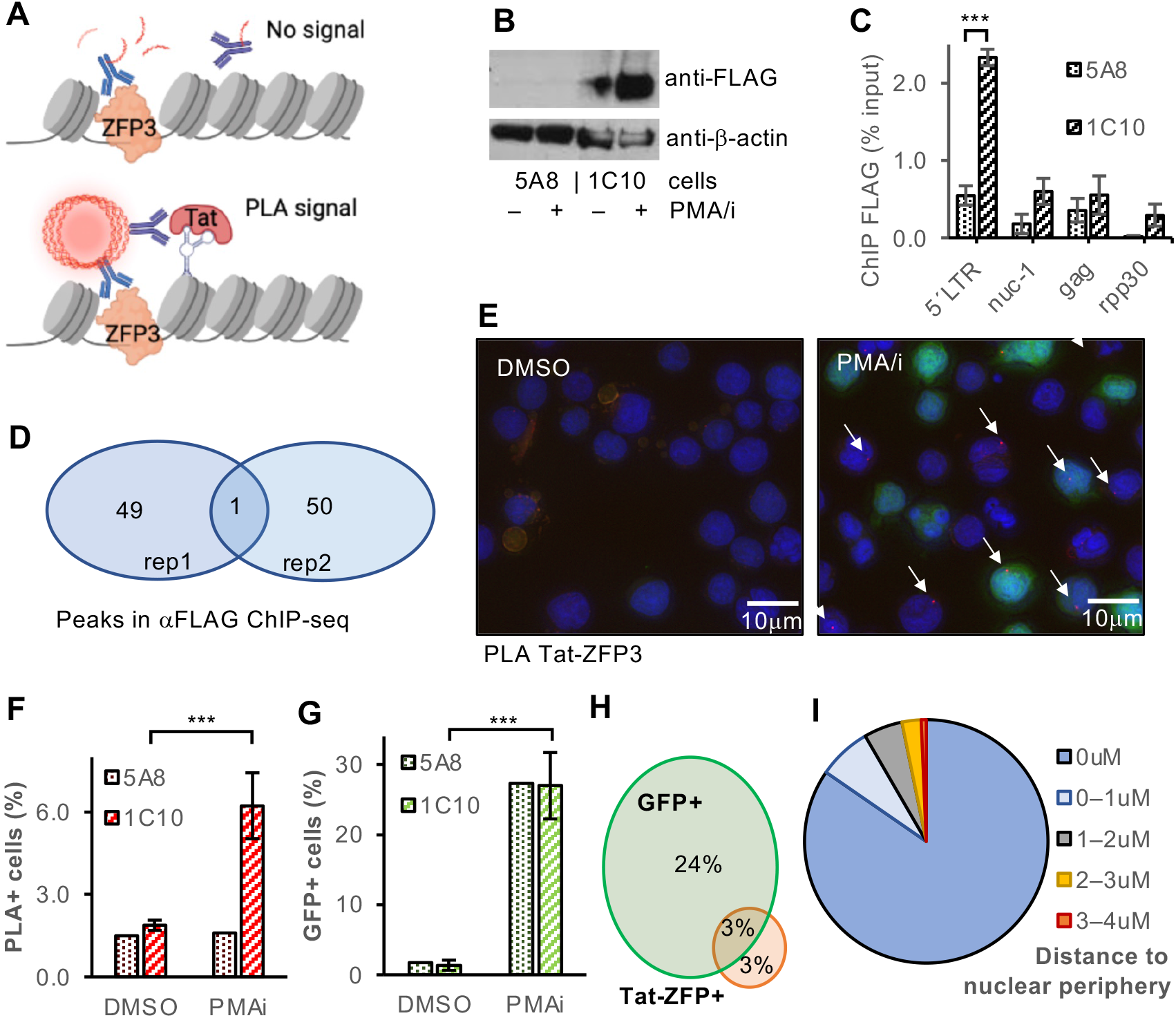
PLA detects Tat at the HIV-1 promoter. (A) The HIV-1 promoter with the zinc finger protein ZFP3 (orange), and an adjacent protein, here Tat (pink) recognized by two oligonucleotide-conjugated antibodies that are ligated to enable rolling circle amplification, and subsequent binding of fluorescently labelled probes during PLA. (B) Immunoblot using antibody against FLAG in 1C10 cells, a clone of HIV-1 containing J-lat 5A8 cells infected with FLAG-tagged ZFP3, and parental 5A8 cells treated with DMSO or PMA/i for 24 h. (C) ChIP-qPCR of FLAG in 1C10 and 5A8 cells (*n*=3, s.e.m.). (D) Venn diagram of MACS2 peaks (filtered: -log(q)>10, size<1kb) from ChIP-seq with anti-FLAG in 1C10 cells (*n*=2). (E) PLA with 1C10 cells treated with DMSO or PMA/i for 16 h. Nuclei were stained with DAPI, green shows GFP expression and red shows PLA spots. White arrows points to the PLA spots. (F–G) Quantification of Tat-ZFP3 PLA^+^ cells (F) and GFP^+^ cells (G), (H) Venn diagram showing the overlap between GFP ^+^ and Tat-ZFP3 PLA^+^ cells, (I) Distance of PLA spots to nuclear periphery (*n*=5). (B, F, G) Student t-test p-values * p<0.05, ** p<0.01, ***p<0.005.

### The engineered zinc-finger protein ZFP3 binds to the HIV-1 promoter

Next, we tested the binding specificity of the engineered ZFP3 to its target DNA sequence. The ZFP3 recognizes a 14nt sequence found in the proviral long terminal repeat (LTR) (position 408–422 at the 5’LTR and 9,493–9,507 at the 3’LTR in the reference HXB2 sequence) which is between the NFκB binding sites and the transcription start site (TSS) (Wang et al., 2014). Chromatin immunoprecipitation (ChIP) followed by qPCR was used with four primer pairs against the 5’LTR, the first transcribed nucleosome *nuc-1*, further downstream *gag* and the human gene *rpp30* as reference. The data confirm that the ZFP3 binds preferentially to the LTR region of the provirus (Fig 1C). The ChIP generated low DNA yield as expected from the low ZFP3-FLAG abundance and high binding specificity. We also performed ChIP followed by massive parallel sequencing (ChIP-seq) to identify other regions in the human genome where the ZFP3 protein might bind. In replicate experiments (*n*=2) we performed MACS2 peak calling (Table S1). Stringent selection criteria (FDR<10^−10^, peak width<1kb in both replicates) identified the HIV-1 provirus as the single binding site of ZFP3 (Fig 1D), at a peak from 293-470 (9,327–9,556) overlapping the expected binding sequence. This confirmed that the ZFP3 protein uniquely marks the HIV-1 provirus in 1C10 cells, which enables the use of the PLA technique to study the microenvironment of the HIV-1 provirus in single cells.

### Detecting HIV-1 activation by proximity of Tat and ZFP3

Our first aim was to detect activation of the HIV-1 provirus by investigating the recruitment of Tat to the HIV-1 promoter. We performed PLA with antibodies against both Tat and the FLAG-epitope of ZFP3 in the 1C10 J-lat cells (Fig 1E). Cells were adhered to coated coverslips and a slightly modified PLA protocol was followed. The low abundance of both Tat and FLAG in the cells prompted us to reduce the background signal of the method. Most notably the concentration of enzymes and antibodies were lowered. As the HIV-1 provirus was linked to the GFP coding sequence, we also measured the GFP expression in the same cells. After completion of the PLA protocol, the native GFP signal was obscured, possibly because of destruction of the GFP 3D-structure. To compensate for this, we used a FITC-conjugated antibody against GFP for detection. Nuclei were counterstained with DAPI. After staining and mounting of the coverslip, the PLA and GFP signals were recorded in a slide scanner scoring 500-5,000 cells for each coverslip. The images were automatically and computationally processed using a custom script. Nuclei were identified and the nuclear area was calculated. In the orange channel for PLA, foci were detected at a range of thresholds for the fluorescent intensity (Fig S1B). Based on control experiments, we calculated an arbitrary PLA threshold, that was used throughout the experiments, but slightly adjusted in few samples to account for experimental variability. The cells with a nucleus containing a single focus were considered PLA^+^. Rare nuclei with multiple foci were not counted. From the green channel, the average GFP intensity over the nucleus was recorded.

In DMSO treated samples, where Tat was not expected to be expressed, the background level of Tat-ZFP3 PLA^+^ cells was 1.9±0.2 % in cells treated with DMSO for 16 h. Upon 16 h of treatment with PMA/i to stimulate the cells and activate the provirus, 5.3±1.2 % (*n*=5) of the cells were PLA positive (Fig 1F). To test the sensitivity of the PLA method, we included the parental 5A8 cells. Tat-ZFP3 PLA nuclear spots were found in 1.5±0.1 % of the 5A8 cells lacking FLAG-ZFP3 regardless of cellular activation status. The activated cells had 27±5 % of GFP^+^ cells in both cell lines, with a background of 1.5±0.3 % in DMSO treated cells (Fig 1G). The unstimulated Tat-ZFP3 PLA^+^ cells contain spontaneously activated proviruses as well as technical artefacts. These cells did not express GFP above background levels (Fig S1C). Flow cytometry data of the 1C10 cell lines showed 0.19 % of unstimulated cells spontaneously cells expressing GFP and 19.3 % of stimulated cells expressed GFP (Fig S1D). The similarity in response to T cell stimulation in the two cell lines differing in ZFP3 status demonstrated that tethering ZFP3 to the HIV-1 promoter did not interfere with the proviral activation. Overall, the Tat-ZFP PLA^+^ and the GFP^+^ cells only partially overlap (Fig 1H). This suggests that the two read-outs capture different though overlapping aspects of the HIV-1 activation.

As the nuclear position of the provirus can be determined in our samples, we calculated the distance from the PLA spot to the nuclear periphery. As expected from literature, most proviruses were at the nuclear envelope or close to the edge of the nucleus (Fig 1I). After T cell stimulation, the provirus was found even closer to the nuclear periphery (Fig S1E)

### Tat is found mainly at the proximity of the HIV-1 promoter in activated cells

Tat under an ectopic promoter has been reported to have affinity for many genomic regions in unstimulated Jurkat cells (Marban et al., 2011; Reeder et al., 2015). This raised our concern as high levels of non-HIV-1 Tat potentially could increase unspecific background signal in our experiments. Therefore, we mapped Tat in our HIV-1 containing 1C10 J-lat cells with and without T cell stimulation. As expected, immunoblotting revealed detectable but low levels of Tat in the unstimulated 1C10 cells. The levels of Tat increased after T cell stimulation (Fig 2A). ChIP-qPCR showed that Tat was preferably found at the HIV-1 promoter in stimulated cells (Fig 2B). To map the genome-wide distribution of Tat in the nucleus, we performed ChIP-seq. Sequence reads were quality controlled to have MAPQ>20. Tat peaks were detected by the MACS2 algorithm where we compared anti-Tat to input for both activated and resting cells. In this experiment with duplicate samples, we identified 78 Tat peaks in both replicates of the stimulated cells and no peaks in unstimulated cells (Table S1). The sequences of the peaks were analyzed for consensus sequence motifs, but none were identified. Among the highly significant regions (FDR<10^−5^ in both replicates), six regions were identified whereof the most significant region was just downstream of the HIV-1 TSS. The other regions included the splice acceptor within the HIV-1 *env* region, as well as four regions of the host genome. We performed a GO analysis of the Tat-associated regions and found no significant terms enriched. The four non-HIV-1 peaks were found in gene-depleted regions with no obvious link to HIV-1 or T cell activation. After PMA/i exposure, a minor local accumulation of Tat was found on chr 2 at the *MAT2A* locus (Fig S1F). This is the genomic integration site of the HIV-1 provirus in these J-lat cells (Chan et al., 2013).

**Fig 2.**
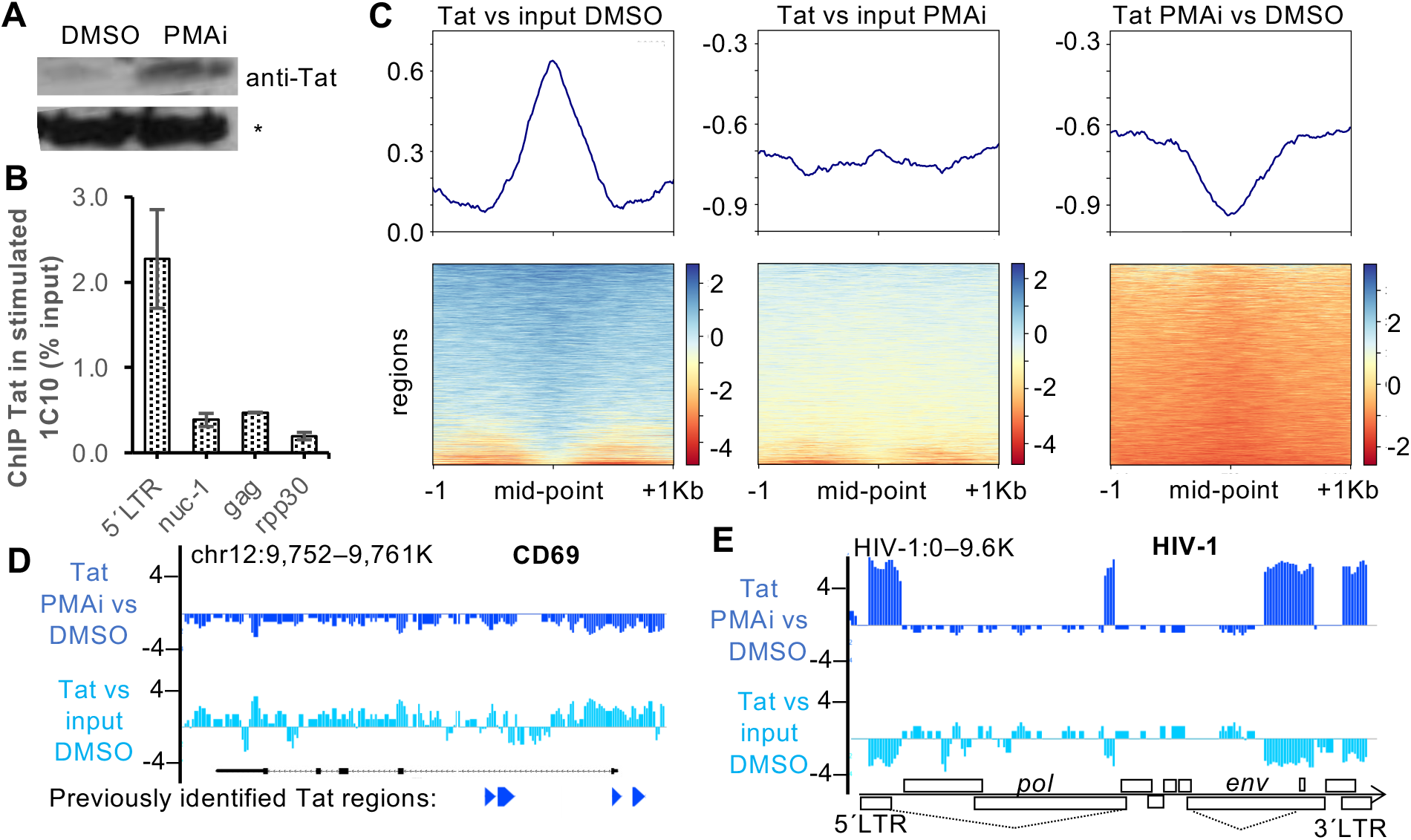
Tat is predominantly found at the HIV-1 promoter in PMA/i activated cells. (A) Immunoblot of Tat. A well-described unspecific band (*) is used as loading control. (B) ChIP-qPCR of Tat at three positions of the HIV-1 provirus and control (*n*=3, s.e.m.). (C) Heatmaps of ChIP-seq against Tat in cells unstimulated (DMSO) or stimulated (PMAi) relative to input or stimulated relative to unstimulated (*n*=2). (D–E) Genome browser view of Tat ChIP-seq, the CD69 locus with the exons in black and the Tat-ChIP peaks from Reeder et al (2015) in blue at the bottom (D) and the HIV-1 locus (E).

Even though we did not clearly identify non-HIV-1 positions associated with Tat binding, we mapped previously identified regions to our dataset. We interrogated the 6,114 peaks associated with Tat binding in unstimulated Jurkat cells (Reeder et al., 2015) (Fig 2C) and indeed observed Tat at these regions in our unstimulated 1C10 cells. However, in the activated cells, the metagene curve was suppressed, with only background variation remaining. The loss of non-HIV-1 Tat during T cell activation became even more noticeable as the decline of Tat at these regions mirrored the Tat profile in unstimulated cells. As an example of these regions, we specifically assessed the locus encoding CD69 (Fig 2D). The browser view demonstrates noticeable levels of Tat at the CD69 promoter locus in unstimulated cells noted previously (Reeder et al., 2015). However, after cellular stimulation, Tat has been redistributed to other regions. This confirms that in stimulated cells under physiological Tat protein levels, Tat binds uniquely to the HIV-1 provirus. Interestingly though, in stimulated cells, apart from the LTR promoter, also an *env* fraction of HIV-1 regions appeared to be associated with Tat, and possibly a fragment within the 3’ *pol* region (Fig 2E).

### PLA and GFP capture different aspects of drug-induced HIV-1 activation

We then sought to determine the effect of latency reversal agents (LRAs) using Tat-ZFP3 PLA. In contrast to other methods that detect transcription irrespective of Tat status, we uniquely detect Tat-dependent transcription. Here we assessed latency reversal 16 h after exposure to six commonly studied LRAs, alone or in combination and at different concentrations. These drugs were the protein kinase C agonists bryostatin and PEP005, histone deacetylase inhibitor romidepsin, BET bromodomain inhibitor JQ1, and the DNMT inhibitor 5-aza-2′-deoxycytidine (5azadC) (Fig S2A). The experiment was controlled by PMA/i-induced T cell activation. We compared the Tat-ZFP3 PLA results to the expression of GFP in the same cells on the same coverslip (Fig S2B). The GFP measurements based on PLA largely recapitulate our previous results from flow cytometry in the parental 5A8 cell line (Lindqvist et al., 2020). T cell stimulation by PMA/i reactivated HIV-1 as detected by both PLA and GFP. However, among the potential LRAs, only the high dose of panabinostat (150 μM) significantly reversed latency detected in both read-outs. Low correlation (0.3) was determined between the GFP and PLA signals. Fitting a straight line through the data points resulted in a goodness-of-fit (R^2^) of 0.1 (Fig S2C). This suggests that different aspects of latency reversal are captured by the two methods, and that the observed GFP signal in the cell model is only partially mediated by Tat-dependent transcription.

### The dynamics of Tat-promoter interactions during cellular activation

To determine the dynamics of Tat recruitment to the HIV-1 promoter during T cell stimulation, we performed a time-series experiment (Fig 3). During 24 h after cellular exposure to PMA/i or DMSO, the appearance of PLA spots representing Tat in the vicinity of the promoter was recorded. As expected, the frequency of cells with a single Tat-ZFP3 PLA spot in the nucleus increased in time in the stimulated cells (Fig 3A). Interestingly, an early background signal could be detected that peaked at 6 h after DMSO addition and then decreased, probably this is an effect of cell handling at the beginning of the experiment. At 15 h after addition of PMA/i, the Tat-ZFP3 PLA spots were above the background of DMSO-treated cells. After 18 h, the frequency of Tat-ZFP3 PLA^+^ cells no longer increased. In contrast, the frequency of GFP^+^ cells steadily increased from 9 h after T cell stimulation (Fig 3B). We also examined the GFP levels in the Tat-ZFP3 PLA^+^ cells. Even though Tat at the HIV-1 promoter was detected in few cells at 12 h after T cell stimulation, these cells were expressing GFP above background (p<0.05) (Fig 3C).

**Fig 3.**
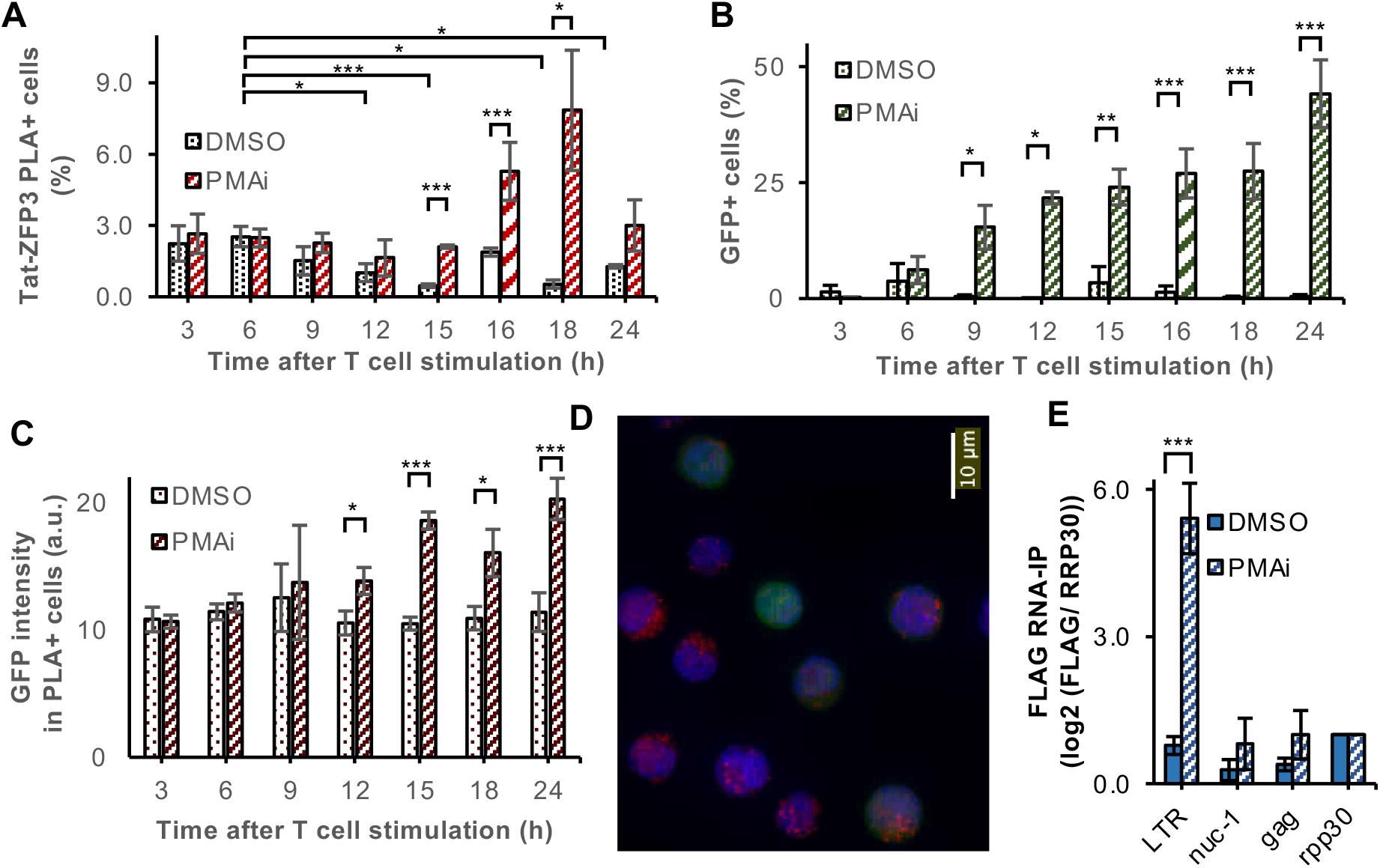
Tat is recruited to the HIV-1 promoter early after T cell activation but binds to cytoplasmic HIV-1 RNA at late timepoints. (A) Time-series with Tat-ZFP3 PLA spots (n=4). (B) Time-series with GFP. (C) GFP intensity of the Tat-ZFP3 PLA+ cells in arbitrary units (D) Micrograph of DAPI-stained (blue) 1C10 cells 24 h after PMA/i exposure. ZFP3-Tat PLA spots in red, GFP in green (E). RNA-immunoprecipitation (RIP) using anti-FLAG. 1C10 cells containing ZFP3-FLAG were treated 24 h with PMA/i or DMSO, qPCR was performed using three primers in HIV-1 and normalized to human *rpp30* (n=3, error bars represent s.e.m.).

### Apart from HIV-1 promoter DNA, ZFP3 also binds to the HIV-1 RNA sequence

At late time points, notably at 24 h after T cell stimulation, we noted irregularities in the PLA data. This was not observed in the unstimulated cells. Upon close examination, we noted a high fraction of nuclei with multiple PLA signals in some of the samples (Fig 3D). PLA spots appear in the cytoplasm and even outside of the cell, and therefore were not recorded as Tat-ZFP3 PLA^+^ cells earlier. We hypothesize that these clusters of PLA signal might be due to ZFP3 binding to the RNA sequence, apart from binding its target sequence in nuclear DNA. The 1C10 cells produce and bud off non-infectious viral particles after activation. Consequently, Tat-ZFP3 PLA signal would also be detectable in the cytoplasm and in viral particles if ZFP3 binds to the LTR in the RNA. To test this hypothesis, we performed RNA-immunoprecipitation (RIP) using an anti-FLAG antibody. The results show that the pulled down FLAG-ZFP3 is associated with RNA from the LTR region of HIV-1 in activated cells (Fig 3E). This signal must originate from the 3’ LTR as the TSS is downstream of the 5’LTR ZFP3 binding region, and thus not present in the HIV-1 RNA. As we are interested in Tat’s role in the initiation of transcription, we decided to perform the following Tat-ZFP3-PLA experiments at 16 h, before viral RNA accumulation in the cytoplasm.

### Activated cells accumulate in G1 while Tat is promoter-proximal in S-G2

Another observation we made here was that, compared to the unstimulated cells, fewer stimulated cells were recovered at the late time points despite same starting material. The 1C10 cells have a population doubling time of 20 h, resulting in the cell numbers more than double in the 24 h following culture addition of DMSO. However, cells grown in media with PMA/i did not change cell numbers in 24 h. This has previously been described as a consequence of a balance between activation-induced proliferation and AICD (Lissy et al., 1998). At 24 h, the growth of the 1C10 J-lat cells in PMA/i containing media was 42±5 % of the DMSO control. In addition, the viability was 83 % in PMA/i treated cells compared to 99 % for DMSO treated cells as evaluated by live-dead stain and flow cytometry (Fig S3A). Together, at 24 h after PMA/i-mediated activation, the surviving fraction was derived from 30 % of the original cells, thus the cell population at this time point did not represent the original population. Given the toxicity of HIV-1 proteins, we expect the surviving fraction to be enriched in cells that have expressed no, low or transient HIV-1 levels.

To further investigate the connection between T cell stimulation and Tat-dependent HIV-1 latency reversal, we interrogated the cell cycle profiles. The nuclear size was used as a proxy for cell cycle stage. In DMSO, the cell cycle profile remains similar throughout the 24 h time course (Fig 4A). After PMA/i treatment, cells appear to arrest in G1 (Fig 4B) as the percentage of cells in late S-G2 goes from 40±3 % at 3 h to 25±4 % at 24 h (Fig 4C). Spontaneous or early PMA/i-induced GFP activation was found predominantly in cells of late S–G2 phases, as expected from HIV-1 latency reversal being linked to metabolically active cells. However, at later time points, GFP^+^ cells accumulate in the G1 phase. With the G1/S arrest activated through the AICD process in the cells, cells with continuous activation of the HIV-1 would be expected to enrich in the G1 fraction, as we observed in the GFP^+^ cells. However, the PLA+ cells displayed a different pattern. At the 15 h time point, before the confounding factor of cytoplasmic spots occurred, 55±6 % of the Tat-ZFP3 PLA^+^ cells were found in the late S–G2 phases. This demonstrates that the cells with Tat at the promoter were not affected by the G1/S arrest (Fig S3B). The lack of accumulation in G1 can only happen if Tat is transiently recruited to the HIV-1 promoter. From this analysis we conclude that whereas GFP detects the dying or arrested cells, Tat-ZFP3 PLA detects the fraction of proliferating cells that will preserve the pool of HIV-1 infected cells post-activation.

**Fig 4.**
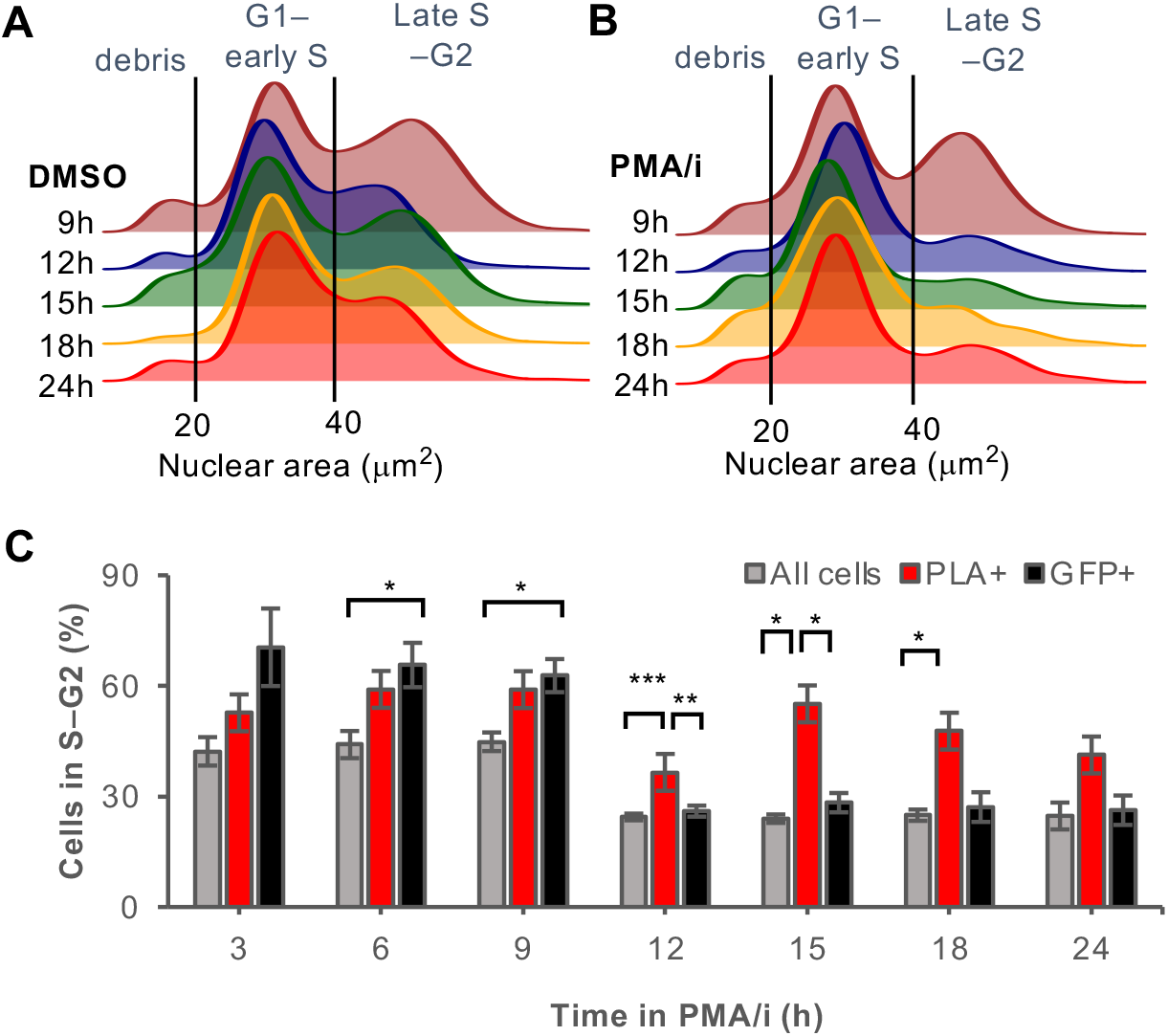
Tat is transcienty bound to HIV-1 promoter in all cell cycle stages. (A–B) Nuclear area was calculated in unstimulated cells (DMSO) (A) or stimulated cells (PMA/i) (B) from the micrographs at 9–24 h chemical exposure. (C) Percentage of cells with nuclear area >40um^2^ corresponding to late S–G2 phase of the cell cycle at different time points after T cell stimulation. All cells in grey, Tat-ZFP3 PLA^+^ cells in red, GFP^+^ cells in black, *n*=5, error bars represent s.e.m., Student t-test p-values * p<0.05, ** p<0.01, ***p<0.005.

### The chromatin composition of the HIV-1 promoter reflects latency reversal potential

Apart from allowing the detection of Tat at the promoter, we can also use the ZFP3 PLA assay to characterize the microenvironment of the HIV-1 provirus in other aspects, *e*.*g*., to determine the suggested heterogeneity of chromatin marks in the reservoir of latently infected cells (Matsuda et al., 2015; Battivelli et al., 2018; Lindqvist et al., 2020). Here we queried the active marks H3K4me1, H3K4me3 and H3K27ac as well as heterochromatin marks H3K9me3 and H3K27me3 in addition to total H3 (Fig 5A). For the first time, we present direct single cell data of chromatin on the reversal of HIV-1 latency. H3 is expected to be detected in all cells. However, we only detect a H3-ZFP3 PLA signal in 6.0±1.8 % of DMSO treated cells and in 11±2 % of cells after PMA/i exposure. This increase most likely reflects a more accessible chromatin after activation. This low recovery prevents comparisons between antibodies, and rather suggests that comparisons should be made between unstimulated and stimulated cells. Strikingly, the activating chromatin marks, H3K4me3, H3K4me1 and H3K27ac seem not affected by stimulation whereas the heterochromatin marks, notably H3K9me3 is detected in more cells after T cell stimulation. H3K9me3 is the only mark that is significantly (p<0.005) different in stimulated compared to unstimulated cells. Worth noting, these data are generated using the surviving fraction of cells that are still intact after 16 h of activation. As we determined previously, T cell stimulation decreases viability and the addition of toxic viral proteins after HIV latency reversal is likely to further reduce viability of virus-producing cells. The fraction of cells present on the coverslips were likely not to have produced large quantities of virus yet. These results are consistent with H3K9me3 at the provirus preventing HIV-1 reactivation and the toxic effects of viral expression.

**Fig 5.**
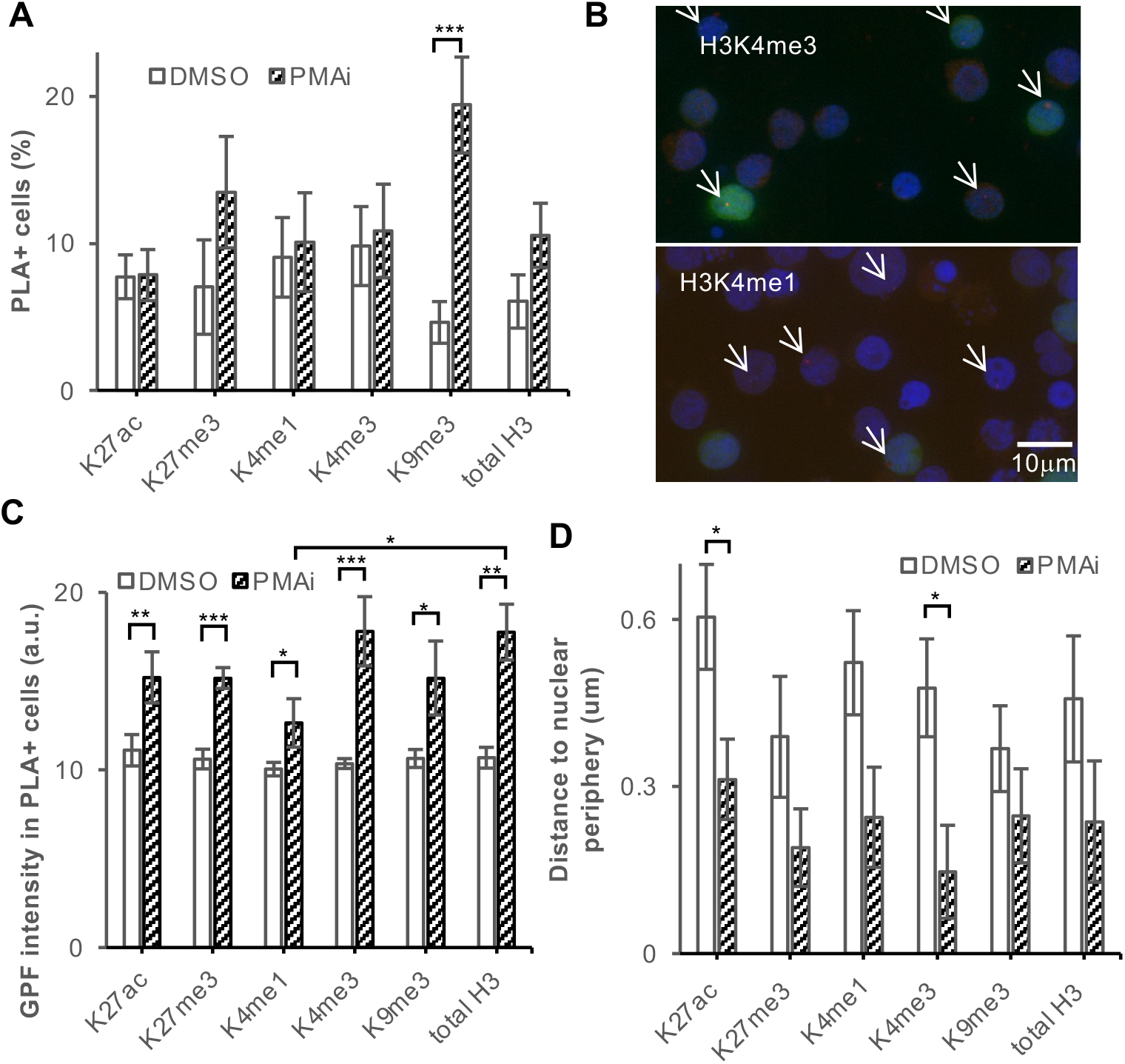
Non-permissive chromatin marks increase in the surviving population after T cell stimulation. (A) H3 modifications found at the HIV-1 promoter at 16 h post T cell stimulation. (B) GFP expression in cells with HIV-1 associated with different marks. (C) Quantification of GFP. (D) Distance to nuclear periphery of the HIV-1 locus. P-values * p<0.05, ** p<0.01, *** p<0.005

Given the nature of the experiment, we had simultaneous access to the GFP expression and PLA signals in the same cells (Fig 5B). To test the reactivation potential associated with the histone modification, we compared the induction of GFP in the ZFP3 PLA^+^ cells after T cell stimulation (Fig 5C). Cells with proviruses associated with all the tested histone modifications had significantly (p<0.05) higher GFP levels after activation. H3K4me3 had the highest relative GFP induction, followed by H3. The weakest evidence for GFP induction was detected with H3K9me3 and the enhancer mark H3K4me1. The GFP level in activated cells was even significantly (p<0.05) lower in H3K4me1-containing proviruses compared to total H3.

We then determined the distances of the HIV-1 proviruses to the nuclear periphery (Fig 5D). Proximity to the nuclear pore facilitates the export of the unspliced RNA required for viral production. The marks of active promoters, H3K4me3 and H3K27ac, were both significantly (p<0.05) closer to the nuclear periphery after T cell stimulation.

### Enhancer H3K27ac is required for Tat-mediated HIV-1 activation

We have previously found that the marks of active enhancers–H3K4me1 in combination with H3K27ac– have a repressive function on HIV-1 (Lindqvist et al., 2020). Employing the ZFP3 PLA tool, we wanted to expand on our previous observation that enhancer-like chromatin modifies the activation status of the HIV-1 provirus. We treated the cells with GNE049, a specific small molecule inhibitor of CBP/P300-mediated H3K27ac at enhancers (Romero et al., 2017; Raisner et al., 2018). As expected, GNE049 exposure led to decreased H3K27ac at the HIV-1 provirus (Fig 6A). However, without H3K27ac, Tat was no longer recruited to the promoter (Fig 6B). HIV-1 was still induced after GNE049 exposure as the GFP levels were induced as in the control T cell stimulation (Fig 6C). This shows that J-lat cells signal HIV-1 latency reversal through GFP despite the lack of Tat at the promoter. However, an enhancer structure of the provirus is required for a Tat-dependent latency reversal of HIV-1.

**Fig 6.**
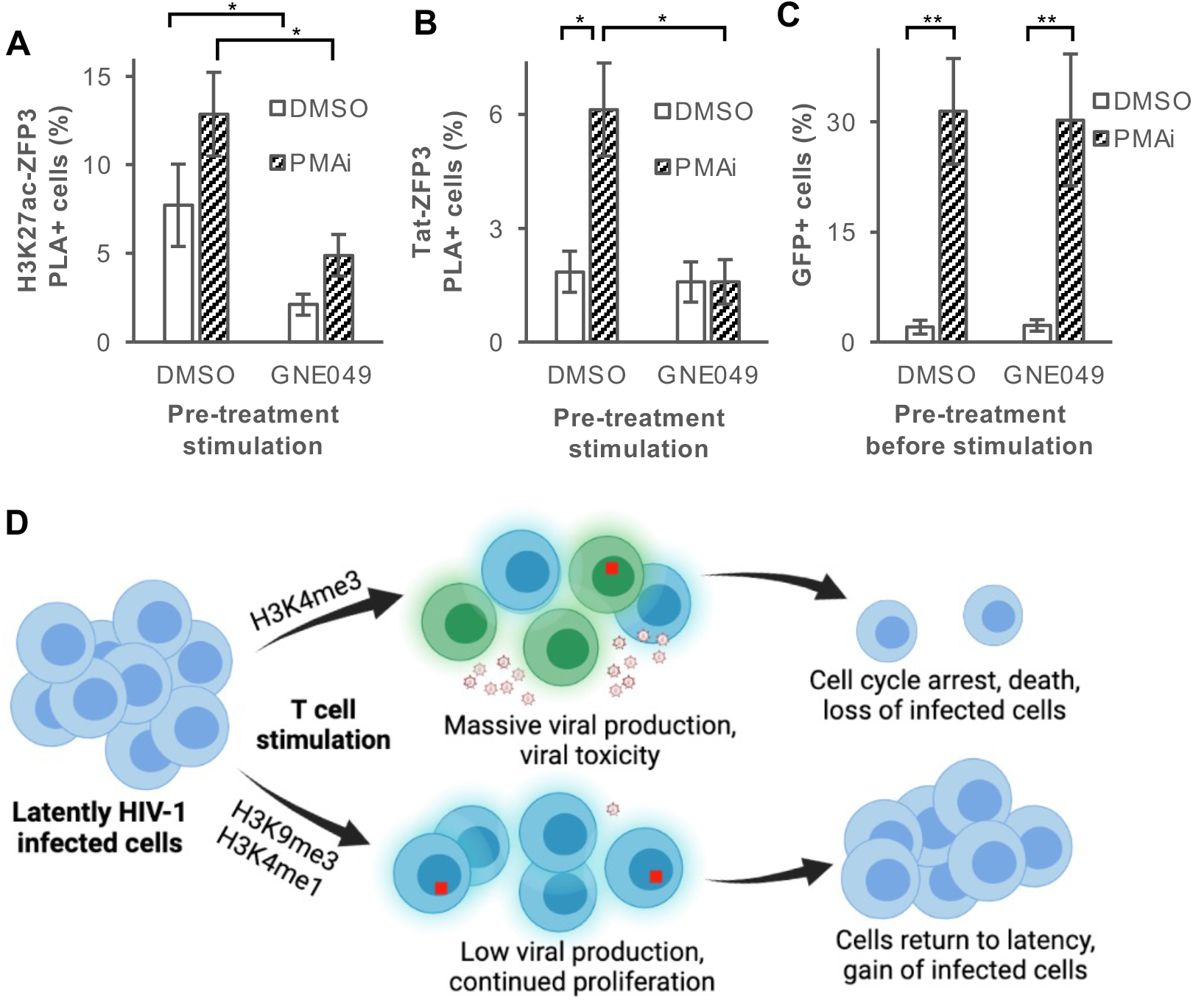
GNE049 reduces H3K27ac and prevents Tat at the promoter without affecting HIV-1 activation. (A) H3K27ac-ZFP3 PLA in cells pretreated with GNE049 (a CBP/P300 inhibitor of enhancer H3K27ac) or DMSO for 3 h followed by 16 h with DMSO or PMA/i. (B) Tat-ZFP3 PLA in the same cells as in (A). (C) Percentage of total GFP^+^ cells treated as in (A). (D) Model of different reservoir compartments after T cell stimulation: upper) high viral production, both dependent and independent of Tat, toxicity leads to cell cycle arrest and cell death; lower) low Tat-mediated viral production in cells with proviral H3K9me3 or H3K4me1, where cells proliferate and return to latency.

## Discussion

Reversing proviral latency in the HIV-1 reservoir is highly complex. Even in a homogeneous population of cells, intercellular differences result in seemingly stochastic proviral reactivation. The processes leading to latency reversal are highly interconnected because of several positive feedback loops, *e*.*g*., Tat regulating its own transcript and HIV-1 inducing cell death in bystander cells that in turn activates HIV-1. To disentangle the different activities, we present a single cell method that uniquely identifies a transient pulse of Tat in an early but essential step during HIV-1 latency reversal. Without Tat enforcing the HIV-1 transcription, cells do not shed viral particles. In combination with simultaneous GFP as a readout of total viral activation we can distinguish productive, Tat-dependent HIV-1 activation from spurious and non-physiological Tat-independent activation. In addition to providing an early readout for HIV-1 latency reversal, the proviral chromatin microenvironment can be interrogated in individual cells. The method is well suited to untangle mechanisms underlying latency reversal. We have here shown that the previously identified enhancer structure at the HIV-1 promoter is required for Tat recruitment and subsequent virus production. Tat initially enables processive HIV-1 transcription, but ensuing acetylation of Tat result in lost affinity for RNA and P-TEFb, consistent with the transient pulse of Tat we observe at the promoter (Kiernan et al., 1999; Kaehlcke et al., 2003). Our observation that ChIP pulls down Tat from within the provirus is intriguing, as the identified regions contain the major splice acceptor sites. The first splice donor is just downstream of the 5’LTR. A spatial structure where the spliceosome physically connects the splice donor and acceptor may explain the apparent non-promoter Tat peaks. This would also time the pulse of Tat to the phase of early mRNA production, in contrast to the later phase of RNAs containing intron-like structures. Proximity to nuclear pores will facilitate nuclear export of transcripts with retained introns when they are bound by viral Rev. Rev disables the Trp-mediated gatekeeper of the nuclear pore (Coyle et al., 2011). As removal of Tat restores HIV-1 latency, the observed transient pulse of Tat allows proliferation of the HIV-1 infected cells and expands the latent reservoir (Razooky et al., 2017).

The HIV-1 reservoir persists through proliferation (Chomont et al., 2009). This proliferation is driven both by antigens and homeostasis (Simonetti et al., 2020). While the decay of the reservoir is slow, the proviral chromatin is dynamic. Even in the absence of activating signals, the cellular fraction with heterochromatin and enhancer chromatin structures at the provirus expands (Lindqvist et al., 2020). We have shown that even in a rather homogeneous cell population, a multitude of different mutually exclusive chromatin structures are present at the HIV-1 integration site. None of the H3 modifications tested here completely prevented HIV-1 reactivation and even proviruses encapsulated in either of the heterochromatin marks H3K27me3 or H3K9me3 produced mRNA after T cell stimulation. H3K27me3 is associated with bistable chromatin and recently it was shown that depletion of PRC1 lead to rapid removal of H2AK119ub. Even though the H3K27me3 mark was still present, transcription is derepressed (Dobrinić et al., 2020). The H3K9me3 mark is a repressive mark but repression depends on the binding of heterochromatin protein 1 (HP1). Different isoforms of HP1 exist. Whereas HP1α and HP1β prevent promoter activity, HP1γ prevents transcription initiation but occupies regions downstream of the promoter in expressed genes, including HIV-1. Elimination of HP1γ has previously been linked to HIV-1 latency reversal (du Chene et al., 2007). The PMA/i-induced GFP in the cells with H3K9me3 at the provirus (Fig 5C) might be explained by the presence of HP1γ.

Based on the data presented here, we propose a model (Fig 6D) of how the proviral chromatin composition affect the latently HIV-1 infected T cells after stimulation. T cell stimulation leads to initial cellular expansion and availability of transcription factors including NFκB. Proviruses present in a chromatin confirmation resembling an active or poised host gene and thus associated with H3K4me3 at the promoter will be transcribed, resulting in mRNA production and protein production. These cells will also induce cell cycle arrest and AICD. In the presence of ART, the new viruses will not be able to infect other cells and this fraction of the latent reservoir will diminish. On the other hand, cells with a different chromatin composition at the provirus, *e*.*g*., H3K9me3 or H3K4me1 will activate the virus to a lesser extent upon T cell stimulus. Thereby these cells will induce apoptosis to a lesser extent. On the contrary, they have the potential to proliferate resulting in an expanded fraction of the reservoir. In this balance, the number of HIV-1 infected cells remain similar, but the HIV-1 reservoir gradually enters a deeper latency. In time, after spontaneous HIV-1 latency reversal and occasional activation of the CD4 cells, this would shape the HIV-1 reservoir, and the initially rare H3K9me3 proviruses becomes more prominent. A progressive reduction of easily activatable proviruses would explain the appearance of post-treatment controllers after long-term ART, despite the number of HIV-1 infected cells remining similar (Namazi et al., 2018). Even though the proviruses in inaccessible H3K9me3 expand, this may not prevent a functional HIV-1 cure (Jiang et al., 2020). However, the more worrisome sub-compartment of the reservoir consists of cells with proviruses embedded in enhancer-like structures. These proviruses are in a stable silent state that remains open and accessible. Short transcripts are continuously produced, yet they are invisible to the immune system. These cells are likely to be responsible for rebound during ART interruptions (Moron-Lopez et al., 2019; Pasternak et al., 2020). Specifically eliminating this subpopulation of cells containing latent but reactivatable HIV-1 will reduce the fraction of the reservoir responsible for rebound viremia. We show here that, by using small molecules to eliminate the enhancer functionality of the provirus, Tat-independent HIV-1 latency reversal potentially eliminates this reservoir fraction.

## Acknowledgements

The authors would like to thank Sara Svensson Akusjärvi for initial work with microscopy. This study was supported by grants from Swedish Research Council (2019-00991), Cancerfonden (12 0412 Pj) and Stiftelsen Läkare mot AIDS Forskningsfond (Fob2020-0004) to JPS, and Center for Innovative Medicine (FoUI-954473) to JPS ans AS. We would like to acknowledge the core facilities MedH Core Flow Cytometry facility (Karolinska Institutet) for providing cell analysis services, and BEA, Bioinformatics and Expression Analysis (Karolinska Institutet) for providing sequencing services.

## Author contributions

BL, WA, JR, LL, TT, JPS performed the experiments. BL, JR and JPS analyzed the results. LL, BBJ, AS, EV provided intellectual input and revised the manuscript, AS and JPS obtained funding, JPS designed the study, wrote the manuscript. All authors read and approved the final manuscript.

## Material and method

### Plasmid construction

The ZFP3 sequence recognizing CGAGCCCTCAGATGC was synthesized by Genescript (NJ, USA). It was cloned into the plasmid pHR-SFFV-KRAB-dCas9-P2A-mCherry by substitution of KRAB-dCas9 cassette using Gibson assembly (New England Biolabs, Cat#E2611). Later the mCherry was substituted with BFP (Gilbert et al., 2014). Primers for Gibson assembly are found in Table S2. pHR-SFFV-KRAB-dCas9-P2A-mCherry was a gift from Jonathan Weissman (Addgene plasmid # 60954 ; http://n2t.net/addgene:60954 ; RRID:Addgene_60954)

### Cell culture

J-lat 5A8 and 1C10 cells were cultured in cytokine-free media (RPMI 1640 medium (Hyclone, Cat# SH30096_01), 10% FBS (Life Technologies, Cat# 10270-106), 1% Glutamax (Life Technologies, Cat# 35050), 1% Penicillin-streptomycin (Life Technologies, Cat# 15140-122)).

### Virus production

pHR-SFFV-ZF3-P2A-BFP, psPAX2 (Addgene, Cat#12260) and pMD2.G (VSV-g) (Addgene, Cat#12259) plasmids were purified with Plasmid Plus Maxi Kit (Qiagen, Cat# 12963). psPAX2 and pMD2.G were gifts from Didier Trono. 293T cells (ATCC, CRL-3216; CVCL_0063) grown in DMEM media (Hyclone, Cat# SH30022_01) were transfected with Lipofectamine LTX with PLUS reagent (ThermoFisher, Cat# 15338100), and after 48 h supernatants were harvested. We determined virus titers (p24) by Lenti-X GoStix Plus (Takara Bio Cat# 631280). 5A8 cells were transduced with 50 ng p24/10^6^ cells. After 3 days, cells plated in 96-well plates for monoclonal cultures. After 14 days, several clones were tested for BFP expression and GFP expression with and without PMA/i exposure.

### Flow cytometry

Cells were stained with LIVE/DEAD Fixable Violet Dead Cell Stain (ThermoFisher, Cat# L34955) and fixed in 2 % formaldehyde for 30 min. Flow analysis was performed on a CytoFLEX S (Beckman Coulter). Individual flow droplets were gated for lymphocytes, viability, and singlets. Data were analyzed by Flowjo 10.1 (Tree Star).

### Proximity ligation assay (PLA)

Cells were washed with PBS and allowed to settle onto poly-l-lysine coated coverglasses (Corning Biocoat Cat#354085). PLA was performed according to the manufacturer’s protocol (Sigma-Aldrich, cat#Duo92007) with a few modifications: PLA plus and minus probes were diluted 1:20, amplification buffer (5×) was used at 10×. Antibodies were used against Tat (Abcam Cat#43014), FLAG M2 (Sigma-Aldrich Cat# F1804 lot SLCF4933), H3 (Abcam, Cat# ab1791), H3K4me1 (Abcam, Cat# ab8895), H3K4me3 (Diagenode, Cat# C15410030), H3K9me3 (Abcam, Cat# ab8898), H3K27me3 (Diagenode, Cat#C15410069), H3K27ac (Abcam, Cat# ab4729). Before DAPI staining and mounting with ProLong Gold Antifade, FITC-conjugated anti-GFP (Abcam, Cat#ab6662) was applied (1:500) for 1 h at ambient temperature protected from light. Slides were sealed with nail polish and stored at 4 °C overnight before imaging. Slides were imaged using a Pannoramic Midi II slide scanner (3DHistech) and images were exported using the CaseViewer application. Images were analyzed with ImageJ (version 2.0.0-rc-69/1.52) and macros developed in house. Each RGB image was separated into separate channels. Based on the blue (DAPI) channel, nuclei were identified. In the red channel, spots (“maxima”) were detected with thresholds 6–48.

### Chemicals to induce proviral activation

Cells were exposed to latency-reversal agents for 16 h. Drugs and chemicals used were phorbol 12-myristate 13-acetate (Sigma-Aldrich, Cat# 79346) final concentration 50 ng/ml, ionomycin (Sigma-Aldrich, Cat# I0634; Lot#106M4015V) final concentration 1 μM, panobinostat (Cayman Chemicals, Cat# CAYM13280) final concentration 30 nM or 150 nM, JQ1 (Cayman Chemicals, Cat#CAYM11187) final concentration 100 nM, bryostatin (Biovision, Cat# BIOV2513) final concentration 10 nM, GNE049 (MedChemExpress, Cat# HY-108435).

### Chromatin immunoprecipitation

ChIP-qPCR was performed using the iDeal ChIP-qPCR protocol (Diagenode, Cat# C01010180). Each ChIP reaction was performed on 2 × 10^6^ cells. Cells were fixed with 1 % formaldehyde for 10 min in room temperature. Sonication was performed at 30 s in eight cycles (Bioruptor Pico, Diagenode, Cat# B01060010). ChIP was performed using anti-FLAG antibody M2 (Sigma-Aldrich Cat# F1804 lot SLCF4933), or anti-Tat (Abcam Cat#43014). ChIP eluates were purified with Wizard SV Gel and PCR clean-up system (Promega, Cat# A9282). Primer sequences are shown in Table S2. PCR reactions were performed with Powerup Sybr green master mix (2×) (ThermoFisher, Cat#A25742) using 40 cycles on an Applied Biosystems 7500 Fast Real-Time PCR System (ThermoFisher).

### Massive parallel sequencing

DNA samples were quantified with Qubit dsDNA HS Assay kit (ThermoFisher, Cat# Q32851) and libraries were prepared using NEBNext Ultra II DNA library kit. Libraries were sequenced on an Illumina Nextseq 550 (75 cycles, single-end sequencing) at the BEA facility (Huddinge, Sweden), according to the manufacturer’s instructions. Raw data from the sequencing reactions (fastq files) were aligned to the hg38 genome assembly with Bowtie2 (version 2.0.6), set to the default parameters. Resulting sam files were converted to bam files using Samtools version 1.4. Bam files were imported into Galaxy (version 21.05, usegalaxy.org) or SeqMonk version 1.47.2.

### Native RNA-IP

2 × 10^7^ cells were activated for 24 h with PMA/i or DMSO. Cells were washed in cold PBS, and KCl, EDTA and RNase Out were added. The cell pellet was lysed (50 mM Tris pH 8,1% TritonX-100,150 mM NaCl, 1 mM DTT, 2 mM orthovanadate, PIC). The cells were rotated in 4 °C for 30 min and centrifugated in 4 °C 13,000 rpm for 20 min. Magnetic beads were washed in TBS-T (20 mM Tris pH 7,4, 150 mM NaCl, 0.05 % Tween20), then in buffer NT2 (50 mM Tris-HCl pH7,4, 150 mM NaCl, 1 mM MgCl_2_, 0.05 % TritonX-100, PIC). For each reaction, 2 μl antibody and 25 μl washed Pierce magnetic protein A/G beads were rotated 1 h in RT, before adding buffer (NT2. 20 mM EDTA, 1 mM DTT, 200 U/ml RNase Out, PIC) and cell lysate. Reactions were rotated overnight at 4 °C. The beads were washed six times. Trizol was added, 5 min RT, followed by addition of chloroform, 5 min. The reactions were centrifuged, and the upper aqueous phase was transferred to a new tube. RNA was purified (RNA clean and concentrator, Zymo research Cat#R1017). RNA, random hexamers 50 nM, dNTP, were incubated 5 min at 65 °C, then on ice 1 min. cDNA was generated using Superscript III (ThermoFisher Scientific Cat#18080093). cDNA was diluted 1:4 and used for qPCR with Powerup Sybr green master mix (2×) (ThermoFisher, Cat#A25742) using 40 cycles on an Applied Biosystems 7500 Fast Real-Time PCR System (ThermoFisher). Primer sequences are found in Table S2.

### Data availability

The ChIP-seq data have been deposited in the GEO database under ID GSE183275. Published dataset from (Reeder et al., 2015) was retrieved from GEO at GSE65689.

## Supplementary material

**Table S1**. MACS2 peaks of ChIP-seq using antibodies against FLAG and Tat after treatment with DMSO or PMA/i for 24 h.

**Table S2**. Primers used for Gibson assembly, ChIP and RIP.

**Fig S1. Detecting HIV-1 using proximity ligation assay (PLA)**.

(A) Flow cytometry of 5A8 and 1C10 cells, showing BFP (V450-A) against SSC (SSC-A). (B) Boxplot with the GFP levels relative to intensity of the PLA spot. (C) GFP intensity in Tat-ZFP3 PLA^+^ cells after 16 h treatment with DMSO or PMA/i. Dotted line shows the background cellular GFP intensity. (D) Flow cytometry of 1C10 cells showing GFP (B525-A) against BFP (V450-A) in unstimulated (DMSO) or stimulated (PMA/i) cells after 24 h (E) Distance between the PLA spot and the nuclear periphery, *n*=4 (F) Genome browser view of *the MAT2A* locus, harboring the HIV-1 provirus.

**Fig S2. PLA and GFP capture slightly different aspects of HIV-1 activation following LRA**.

(A–B) Response to latency reversal agents (LRAs) in J-lat 1C10 detected by PLA Tat-ZFP3 (A) and GFP (B). (C) Correlation between GFP and PLA spot. *n*=5, error bars show s.e.m.

**Fig S3. Cells accumulate in G1–early S following PMA/i-mediated stimulation**.

(A) Percentage of live 5A8 cells after T cell stimulation recorded by live/dead cell stain using flow cytometry (*n*=3). (B) Percentage of 1C10 cells with nuclear area 20–40 μm^2^ corresponding to G1–early S phase of the cell cycle. All cells in white, Tat-ZFP3 PLA^+^ cells in red, GFP^+^ cells in green, *n*=5 error bars represent s.e.m., Student t-test p-values * p<0.05.

## Graphical abstract

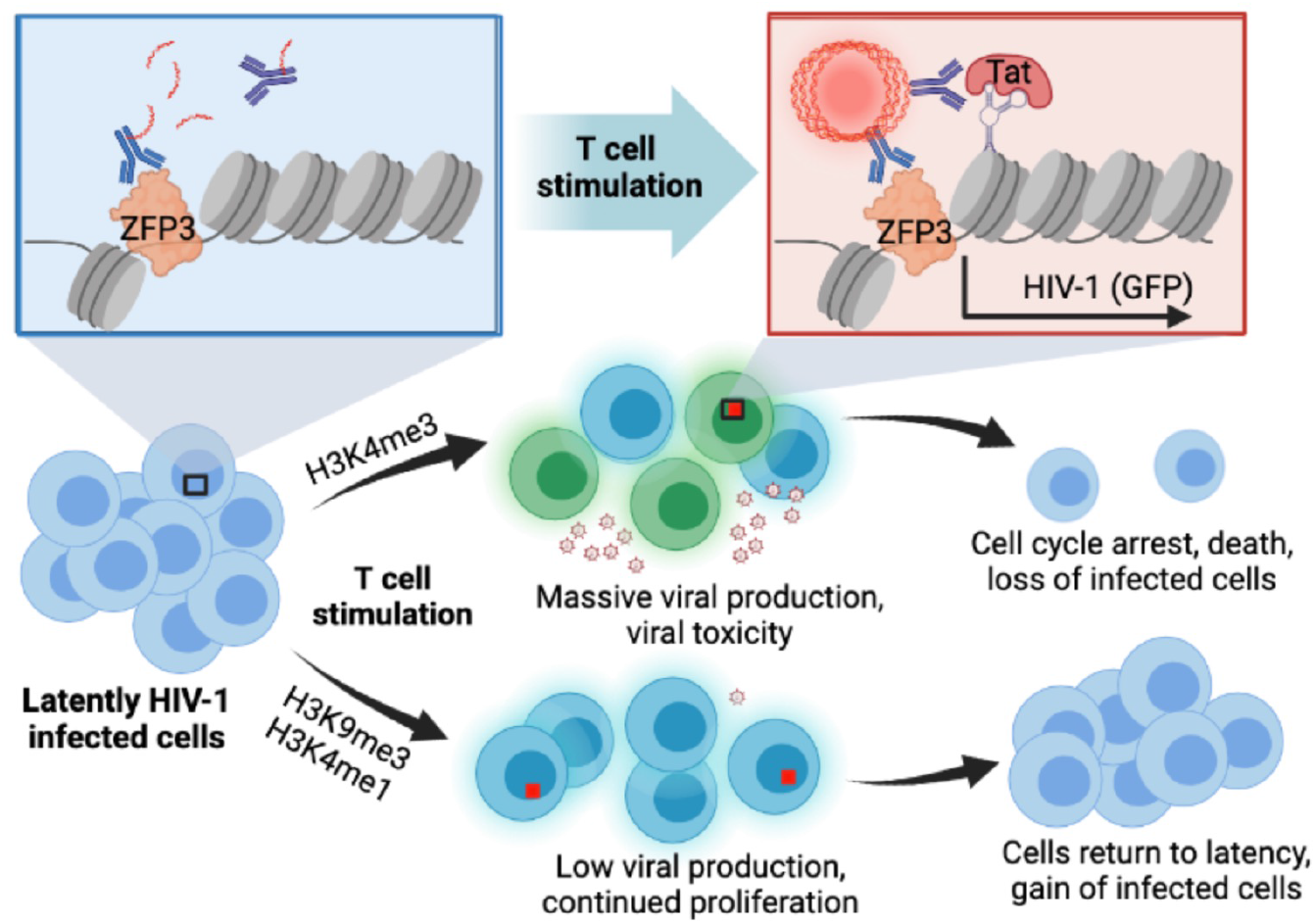

