## Supplementary figures and images for "T cell stimulation remodels the latently HIV-1 infected cell population by differential activation of proviral chromatin"

### Supplemental figures

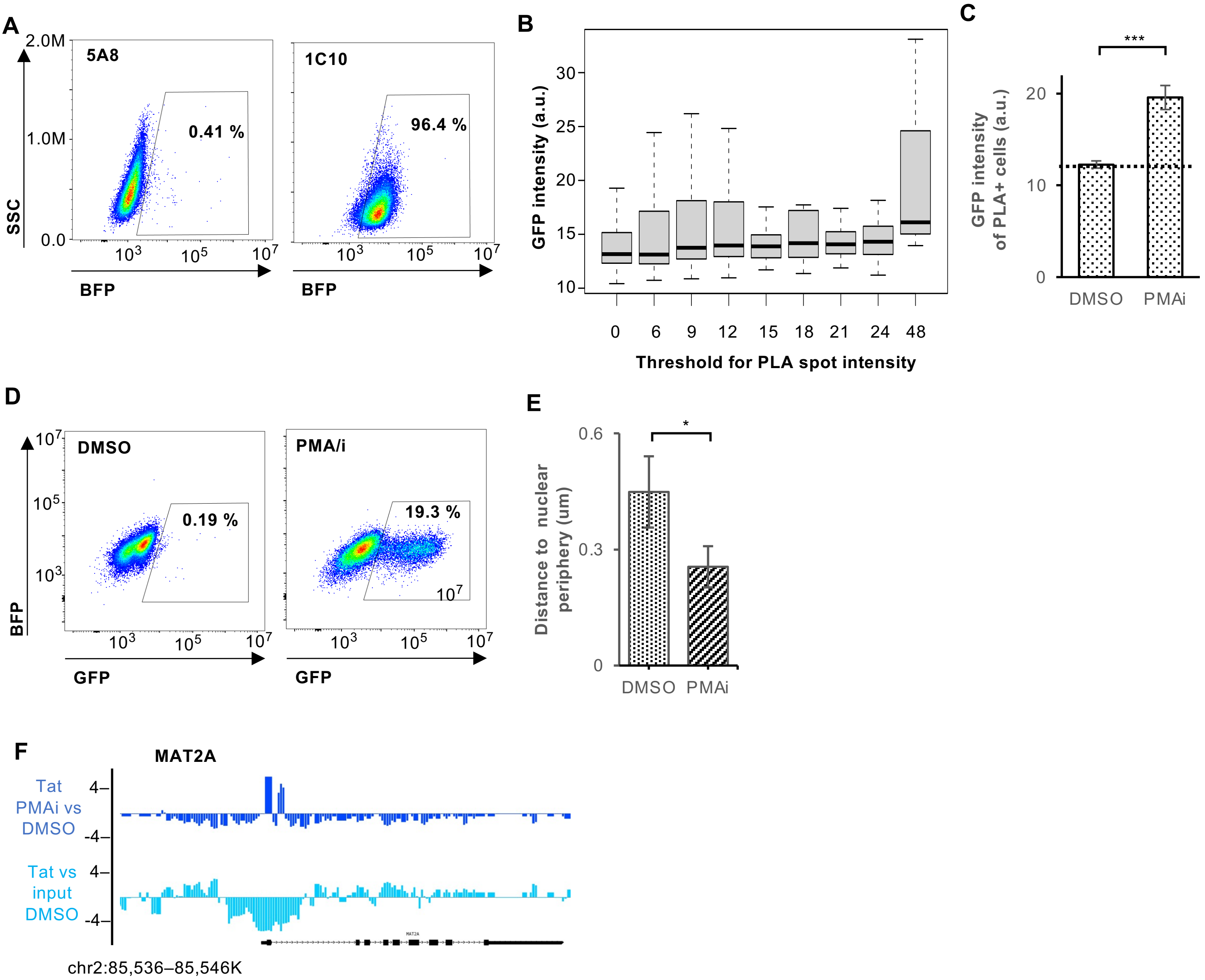

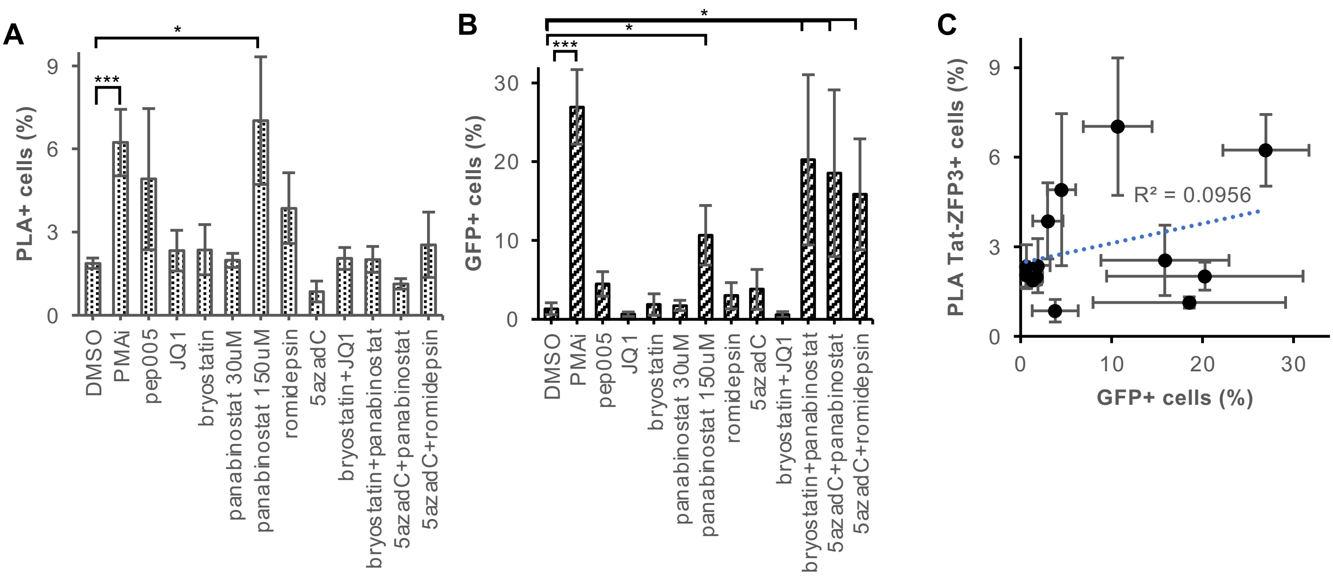

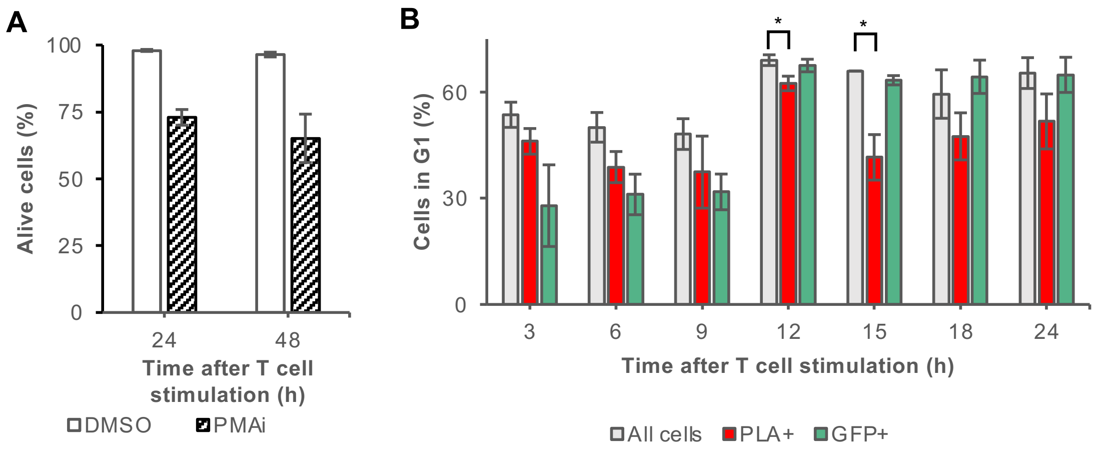
